# Metabolite accumulation mediates the shift between the High Oxygen Consumption and Low Oxygen Consumption phases in the Yeast Metabolic Cycle

**DOI:** 10.64898/2025.12.01.691504

**Authors:** Salvador Casani-Galdon, Shidong Xi, Jane Mellor, Sonia Tarazona, Ana Conesa

## Abstract

The Yeast Metabolic Cycle (YMC) is a molecular system that serves as a model to study the internal clock that maintains homeostasis in complex organisms. Traditionally, this ultradian rhythm has been studied in the three phases where mature mRNA transcripts show peak accumulation. However, recent studies have shown that the YMC can be interpreted as a two-phase cycle based on altered redox states, known as the high (HOC) and low oxygen consumption (LOC) phases. The length of the HOC phase is fixed and its frequency is nutrient dependent but the nature of the HOC to LOC transition is poorly defined. Here, we use multivariate statistics to integrate metabolic, chromatin and transcriptional changes across the YMC to study the levels of organization that connect them. Our model reveals that both the HOC-LOC and LOC-HOC phase transitions in the YMC are coordinated by accumulating metabolites, reflecting cellular energetics and redox state. We propose that the cycling behavior of chromatin states, transcription and transcripts is a consequence of accumulating metabolites at phase transitions, which function by modulating protein activity and coordinating biochemical pathways to maintain cellular homeostasis.

## Introduction

Understanding the intricate patterns of regulatory interactions across molecular layers is a central objective in systems biology^1,2^. Epigenetic modifications and chromatin dynamics are proposed to orchestrate gene expression, triggering signaling cascades that control cellular functions and metabolic circuits. Metabolism, in turn, not only governs cellular states but also contributes to the regulation of epigenomic landscapes^3^. Despite these known relationships, the precise interconnections across these molecular layers remain poorly understood, particularly regarding the specificity of metabolic control over gene expression^4^.

One biological scenario where this inter-regulation is of paramount importance is in ultradian rhythms, where tightly controlled feedback loops between chromatin dynamics, gene expression, and metabolic processes are proposed to establish cyclic patterns of cellular state^1,5^. The Yeast Metabolic Cycle (YMC) is a well-characterized ultradian rhythm in which metabolites oscillate under continuous nutrient-limiting conditions, and the cycling of epigenetic marks has been shown to correlate with mRNA transcript levels^6^. In the YMC, mRNA-sequencing data reveal a three-phase cycle: Oxidative (OX), Reductive Building (RB), and Reductive Charging (RC) phases (Figure 1A)^7^. Subsequent studies have explored the cycling of epigenetic marks and metabolites in relation to nascent transcription and mature mRNA transcripts, suggesting that although transcripts cycle in three phases, nascent transcription may be regulated in only two, pointing to the complexity of epigenetic regulation^8^.

**Figure 1.**
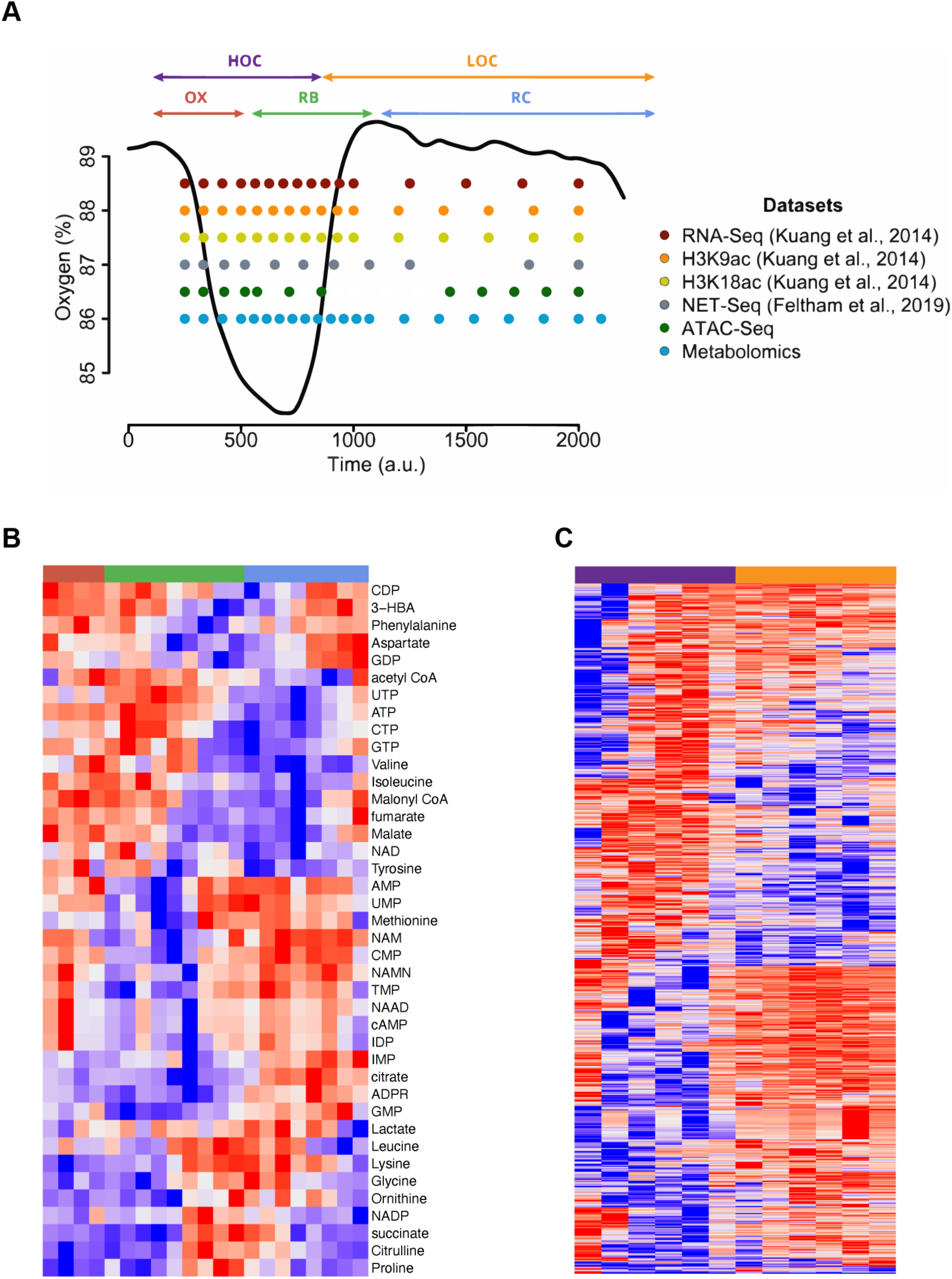
Multi-omic dataset in the Yeast Metabolic Cycle. (A) Schematic of sampling points across the Yeast Metabolic Cycle (YMC) for the six datasets that are integrated in this study. (B) Heatmap of the differentially abundant metabolites across the YMC. (C) Heatmap of the differentially abundant gene promoters across the ATAC sampling points.

While numerous studies have explored multi-layered dynamics in the YMC and similar systems^9^, there is still a lack of generalizable frameworks that integrate chromatin, transcription, and metabolism to generate hypotheses about regulatory interactions and their connection to cellular states. Here, a general framework for integrative, multiscale modeling of the chromatin-transcription-metabolism axis is presented. Using the YMC as a model system, multivariate methods are leveraged to explore regulation across these molecular layers. This approach includes Partial Least Squares Path Modeling (PLS-PM), which allows the evaluation of plausible models that connect multi-modal cellular states to the molecular features of these complex, multi-layered interactions.

## Results

### High-resolution sampling in the Yeast Metabolic Cycle

The Yeast Metabolic Cycle (YMC) is an extensively characterized ultradian rhythm, with available datasets on RNA-seq^6^, NET-seq^8^, and histone marks such as H3K9ac^6^ and H3K18ac^9^, collected from multiple samples throughout the cycle (**Figure 1A**). Although other studies have characterized the fluctuation in metabolites and chromatin accessibility^10,11^, the sampling was done at a lower resolution compared to the transcriptomics and histone modification datasets^6,9^. Our goal was to create a multilayered, multi-omics dataset by complementing existing data with high-resolution metabolomics and ATAC-seq, thereby enabling a comprehensive exploration of metabolism and chromatin accessibility.

We sampled the cycle in quintuplicate to investigate the impact of metabolic fluctuations in the Yeast Metabolic Cycle (YMC). A total of 21 samples were collected across the YMC in each sampling experiment (**Figure S1A**), measuring a total of 63 metabolites (**Table S1**).

Metabolomic profiling revealed temporal oscillations of individual metabolites, as previously described^11^. The five technical replicates captured the progression of the YMC (**Figure S1A-B**) and demonstrated strong agreement across the five sampled cycles (**Figure S1C**). We combined the five replicates across the 21 time points (**Figure 1A, S1B**), discarding one outlier for each sampling point to enhance data quality. The high resolution of metabolic sampling revealed distinct groups of metabolites coordinating across the cycle, such as NTPs with acyl-CoAs, and NMPs with NAMN and ADPR (**Figure 1B**).

Previous studies have hypothesized that metabolic fluctuations impact gene regulation in the YMC through chromatin dynamics^6^. To assess chromatin accessibility at gene promoters in the YMC, we performed Assay for Transposase-Accessible Chromatin with sequencing (ATAC-Seq)^12^ at 13 sampling points across the cycle (**Figure 1A**). We observed a distinct pattern of promoter accessibility along the YMC (**Figure 1C**), with a sequential order that mimics the patterns seen in metabolites (**Figure 1B**). Some genes exhibited open chromatin in their promoters early in the cycle, while others became accessible at later stages. Next, we develop a model to explore the precise temporal relationships between chromatin dynamics, metabolite oscillations and gene expression in the YMC using the six high resolution datasets (**Figure 1A**).

### Metabolites, chromatin and mRNA transcripts follow distinct regulatory dynamics in the Yeast Metabolic Cycle

Previous studies have suggested that the YMC follows a three-phase scheme consisting of an Oxidative phase (OX), followed by a Reductive Building phase (RB) and a Reductive Charging phase (RC)^6,7^, while other authors suggest only two distinctive phases: High Oxygen (HOC) and Low Oxygen (LOC) Consumption phases^1^. We evaluated how the different omics datasets presented in this study fit into these two schemes. For each dataset, we clustered features using k-means and calculated the optimal number of clusters for each omics dataset by evaluating 20 different metrics of clustering quality. Based on these metrics, we selected the most frequently endorsed number of clusters for each omics dataset (**Figure S2A**).

Clustering of RNA-seq data supported the three-phase scheme, with 1,162 transcripts peaking at OX, 653 in RB and 1,278 in the RC phase (**Table S2**). Analysis of the genes encoding transcripts peaking at each phase confirmed previous observation that mature transcripts peak in three distinct phases (**Figure 2A**, **Figure S2A**). In contrast, clustering of the nascent transcripts (NET-seq), H3K9ac and H3K18ac ChIP-seq, and ATAC-seq data supported a two-phase scheme (**Figure 2A**, **Figure S2A**). All four datasets showed an even number of features distributed between the HOC and LOC phases (**Table S2**). These findings suggest that transcriptional regulation in the YMC follows two-phase dynamics that encompasses chromatin accessibility, histone modifications and nascent transcription, supporting the HOC/LOC scheme and that mRNA transcripts in the RB phase are subject to additional post-transcriptional regulation, creating the third cycling phase^8^. The two histone modifications showed high correlation in their promoter signals (**Figure S2B**).

**Figure 2.**
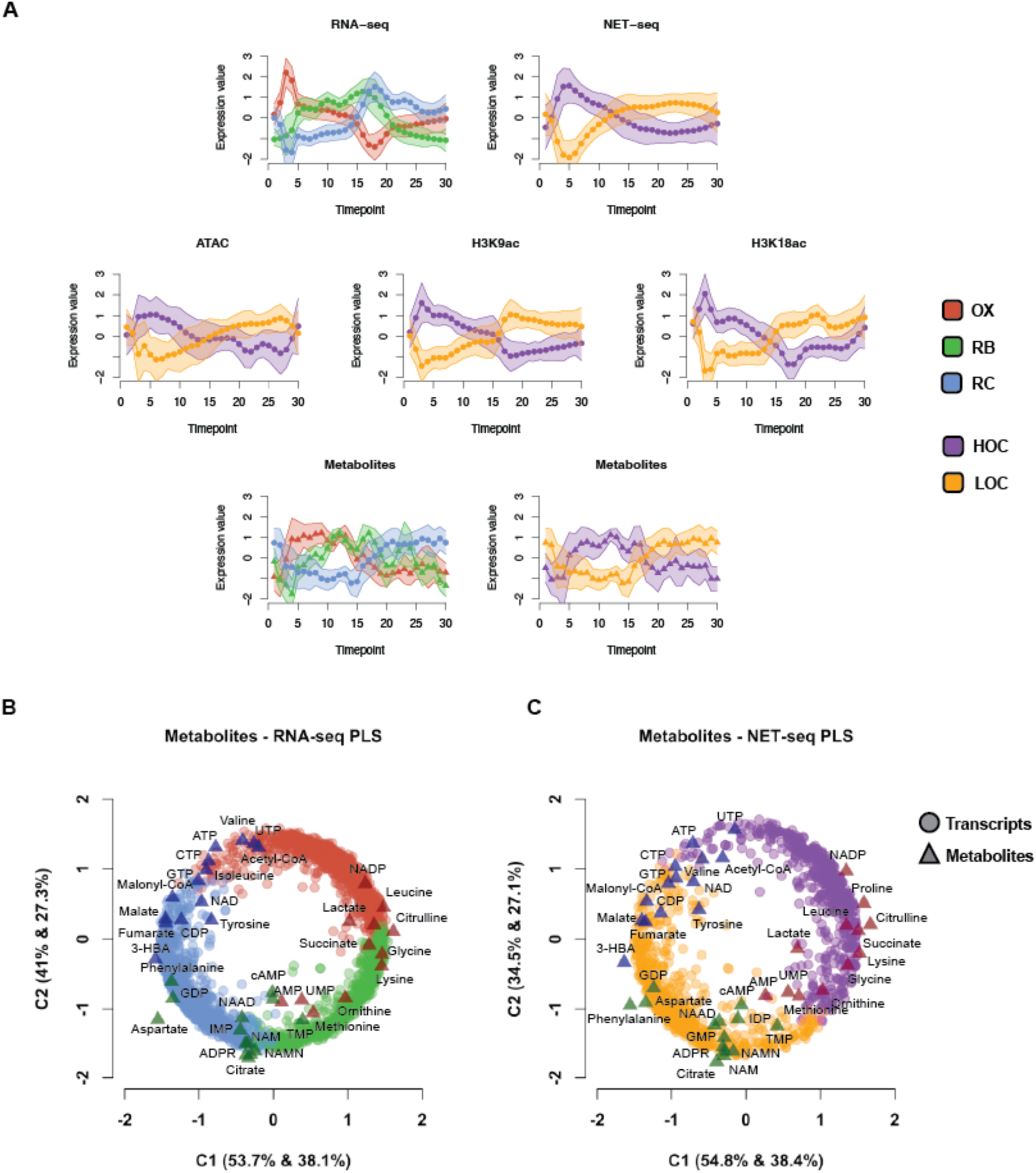
Metabolites peak between transcript phases in the YMC. (A) Cluster profiles for each molecular type analyzed in this study. Transcripts (top) were modeled in three phases for RNA-seq and two phases for NET-seq, chromatin accessibility measurements (middle) were clustered in two phases. Metabolites (bottom) could be clustered in either two or three clusters. (B) Partial Least Squares Modeling of mature transcripts (Y) and metabolites (X) (C) Partial Least Squares Regression of nascent transcripts (Y) and metabolites (X).

Uniquely, metabolites could be classified into both two- and three-phase schemes, with an equal number of indices in each case (**Figure 2A**, **Figure S2A**). The distinct phase distribution of metabolites is striking, perhaps reflecting their involvement in both chromatin state regulation (two phase scheme), and reflecting enzyme/protein activity (three phase scheme).

### Metabolite fluctuation reflects the transcriptional state of the cycle

To resolve this dichotomy in metabolite phasing, we closely examined their dynamic fluctuations over time and observed patterns of accumulation between both mRNA and nascent transcript phases (**Figure 2A**). To explore these relationships in more detail, we analyzed their shared dynamics using Partial Least Squares regression (PLS)^13^ models. PLS regression is a dimension reduction technique that maximizes covariance between the latent variables in the explanatory matrix (***X***) and the latent variables in the response matrix (***Y***). Here we used the metabolites as the explanatory matrix (***X***) and modeled RNA-seq (**Figure 2B**) or NET-seq data (**Figure 2C**) as the response matrix (***Y***).

The joint analysis of RNA-seq and metabolites through PLS regression revealed two major components of covariation that explained 94.7% and 65.4% of the variance in mature transcripts and metabolites, respectively (**Figure 2B**). This integrated analysis reinforced the three-phased cycling of metabolites when analyzed jointly with mature transcripts. Different groups of metabolites corresponding to the three clusters (**Table S2**), which peaked at three points of the cycle, projected in the common space between the mature transcript phases (**Figure 2B**). This suggests that distinct metabolites accumulate either upstream or downstream of regulatory events controlling mRNA synthesis and degradation, and thus net levels of mature mRNA in the cycle. Notably, acetyl-CoA accumulated at the beginning of the OX phase, where it has been reported to mediate histone acetylation and drive entry into OX phase^14^. Furthermore, mRNAs from genes involved in fatty acid degradation show peak levels at the end of the RC phase, which is consistent with the accumulation of acetyl-CoA, NTPs and NDPs during the transition from RC to OX. At this point in the YMC, cells are likely to be in a high energy state, enabling cells to grow and a small proportion (<10%) to enter the cell division cycle^8,15^. The metabolites that accumulated between the OX and RB mature mRNA phases were primarily amino acids, possibly linked to cells entering cell division^16^.

Conversely, metabolites associated with low redox potential, such as citrate, NMPs and NAD-derivatives (such as NAAD or NMN)^17^, accumulated at the beginning of the RC phase. Their peaks are preceded by the accumulation of mRNA transcripts from mitochondrial genes during the high energy state RB phase. We hypothesized that the high energy requirements of the RB phase may lead to the accumulation of low-energy metabolites, signaling the transition into the RC phase.

We then applied the same principles to jointly analyze the NET-seq and metabolomics datasets (**Figure 2C**). PLS regression revealed two major components of covariation that explained 89.3% and 65.5% of the variance in nascent transcripts and metabolites, respectively (**Figure 2C**). Like the three-phase relationship between metabolites and transcripts, metabolites also accumulate at the transitions between the HOC and LOC phases. However, there is one major difference; amino acids accumulate in the middle of the NET-seq HOC phase without leading to a transition into a new phase of nascent transcription and despite amino acids being reported to affect the epigenetic landscape^18^. The accumulation of amino acids is likely to reflect the high energy state metabolism of this phase.

From this analysis we hypothesize that metabolites may act as signaling messengers transmitting signals in the form of chromatin changes that drive nascent transcription in the HOC and LOC phases, or may reflect the accumulation of byproducts of metabolism at the transition points between the three phases, as the cells cycle between different states.

### HOC and LOC YMC phases are enriched in metabolite-sensitive TF-binding sites

Earlier studies characterized chromatin dynamics in the YMC in the three phases described for RNA-seq^6^. Subsequent studies propose that only mature transcripts peaking in the OX and RC phases are regulated by cycling nascent transcription^8,9^ and predict that chromatin dynamics would reflect the onset of transcription. To assess the impact of changes in chromatin opening on changes in nascent transcription, we computed the differential ATAC-seq peaks over the HOC and LOC sampling points in the YMC (**Figure 3A**). We obtained a total of 154 differential peaks, of which 64 were higher in the LOC phase and 90 were higher in the HOC phase. We studied the TF motifs enriched in either group. Interestingly the HOC motifs include glucose response TFs such as Azf1 or Sko1 (**Figure 3B**). The LOC peaks include motifs from TFs involved in oxidative stress response such as Rox1 or Stb5 (**Figure 3C**). These results indicate that the chromatin accessibility recapitulates the biochemical states defined for HOC and LOC phases.

**Figure 3.**
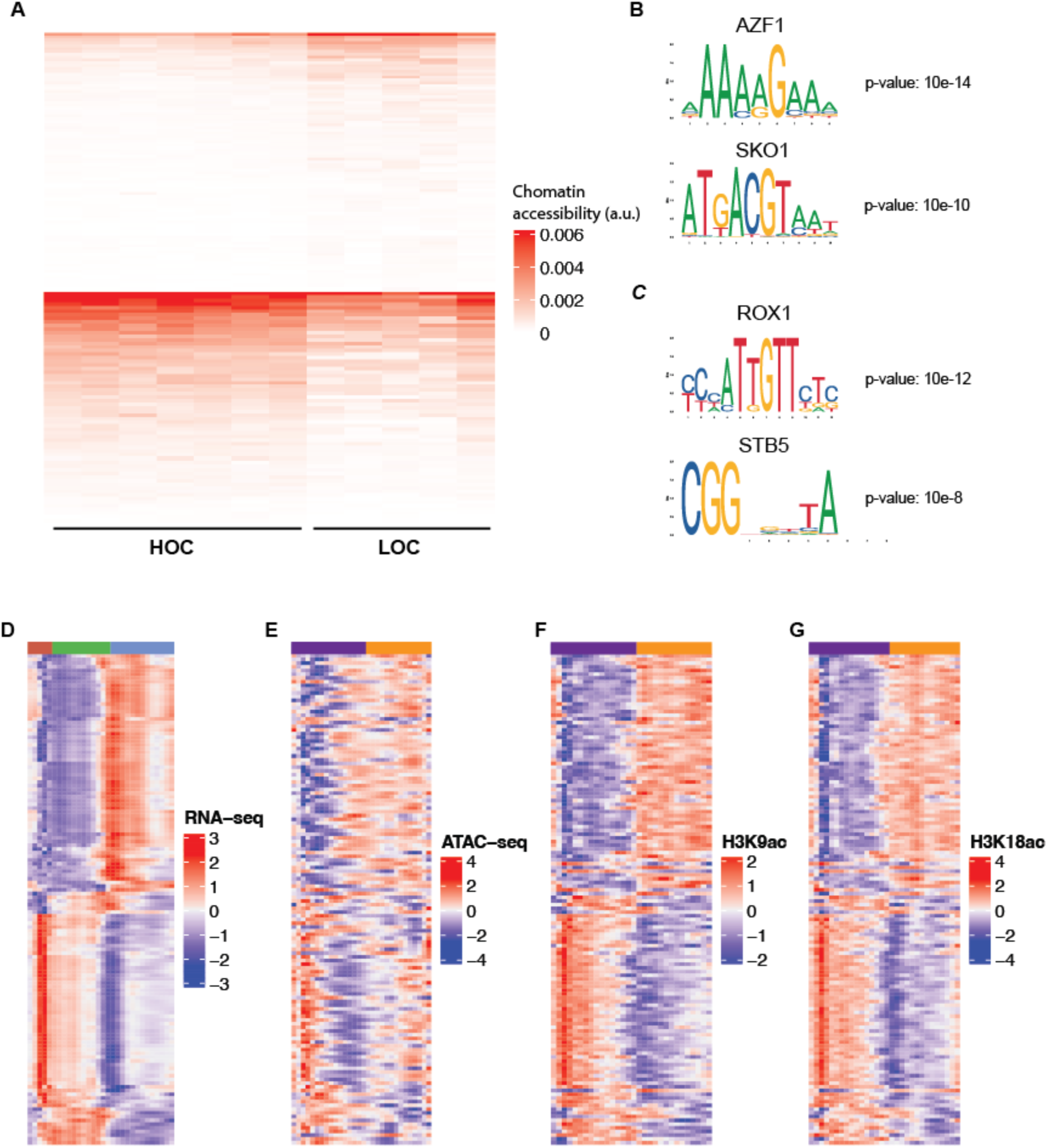
Chromatin accessibility cycles resemble OX and RC mRNA phases. (A) Accessibility heatmap of ATAC-seq regions differentially accessible between HOC (bottom) and LOC (top). (B) Transcription factor binding site motifs associated with HOC ATAC-Seq peaks. (C) Transcription factor binding site motifs associated with LOC ATAC-Seq peaks. Profiles of common genes between (D) RNA-seq (E) ATAC (F) H3K9ac (G) H3K18ac

### Chromatin oscillations coordinate with mRNA transcripts in the OX and RC YMC phases

Next, we examined which genes associated with differentially expressed mature transcripts (RNA-seq; **Figure 3D**) also exhibit significant changes in ATAC-seq (**Figure 3E**) and histone modification levels (H3K9ac and H3K18ac; **Figure 3F&G**). Of the 132 genes encoding mRNAs in this class, 41% peaked in OX and 49% peaked at RC. Only 14 (10%) of these genes were assigned to the RNA-seq RB phase, supporting the notion of the genes expressed in this phase not being subjected to major chromatin regulation. In addition, chromatin dynamics reflects the cycling of the mRNA transcripts. H3K18ac and H3K9ac show a strong correlation with RNA-seq data for these genes (*r* = 0.72 and 0.73, respectively, p-values < 2.2e-16), whereas ATAC measurements exhibit a lower correlation, likely due to higher noise levels in this type of data (*r* = 0.36, p-value < 2.2e-16). In summary, the molecular data capturing chromatin changes showed high correlation with transcript changes in the OX and RC phases but not with those in the RB phase. This suggests that the physicochemical properties defining the HOC and LOC phases lead to dynamic chromatin opening, reflecting the onset of transcription in two phases of the YMC.

Next, we set out to generate an integrative model to assess how metabolic fluctuations influence transcription through chromatin dynamics, and how these regulatory signals mediate transitions between the HOC and LOC phases.

### Modeling molecular relationships across multiple omics into a single model

Given the distinct cycling patterns of the molecular features along the YMC, and the sequential interplay between chromatin dynamics, nascent and mature transcript levels, and metabolite concentrations, we sought to model their relationships and uncover potential regulatory crosstalk. Partial Least Squares Path Modeling (PLS-PM) is a multivariate technique that models relationships among observed and latent variables^19^. We hypothesize that molecules from one phase can influence the regulation of molecules in the subsequent phase (e.g. metabolites in RC affecting histone acetylation in HOC). Accordingly, we developed a PLS-PM framework for the YMC to explore of these relationships. This analytical framework comprises two components: the inner and outer models^20^. The inner or structural model represents a theoretical network describing the connections between latent variables (LV), unmeasured concepts that, in our case, correspond to the cycling phases of each molecular layer in the YMC. We defined a model connecting these latent variables based on the theoretical molecular regulatory flow within the YMC, testing the hypothesis that certain metabolites trigger chromatin changes that in turn affect gene expression during the LOC-HOC and HOC-LOC phase transitions (**Figure S3A**). The outer (or measurement) model defines the measured variables (omic features in our case) that are proposed to contribute to each latent variable (illustrated in **Figure 4A** for early RC metabolites).

**Figure 4.**
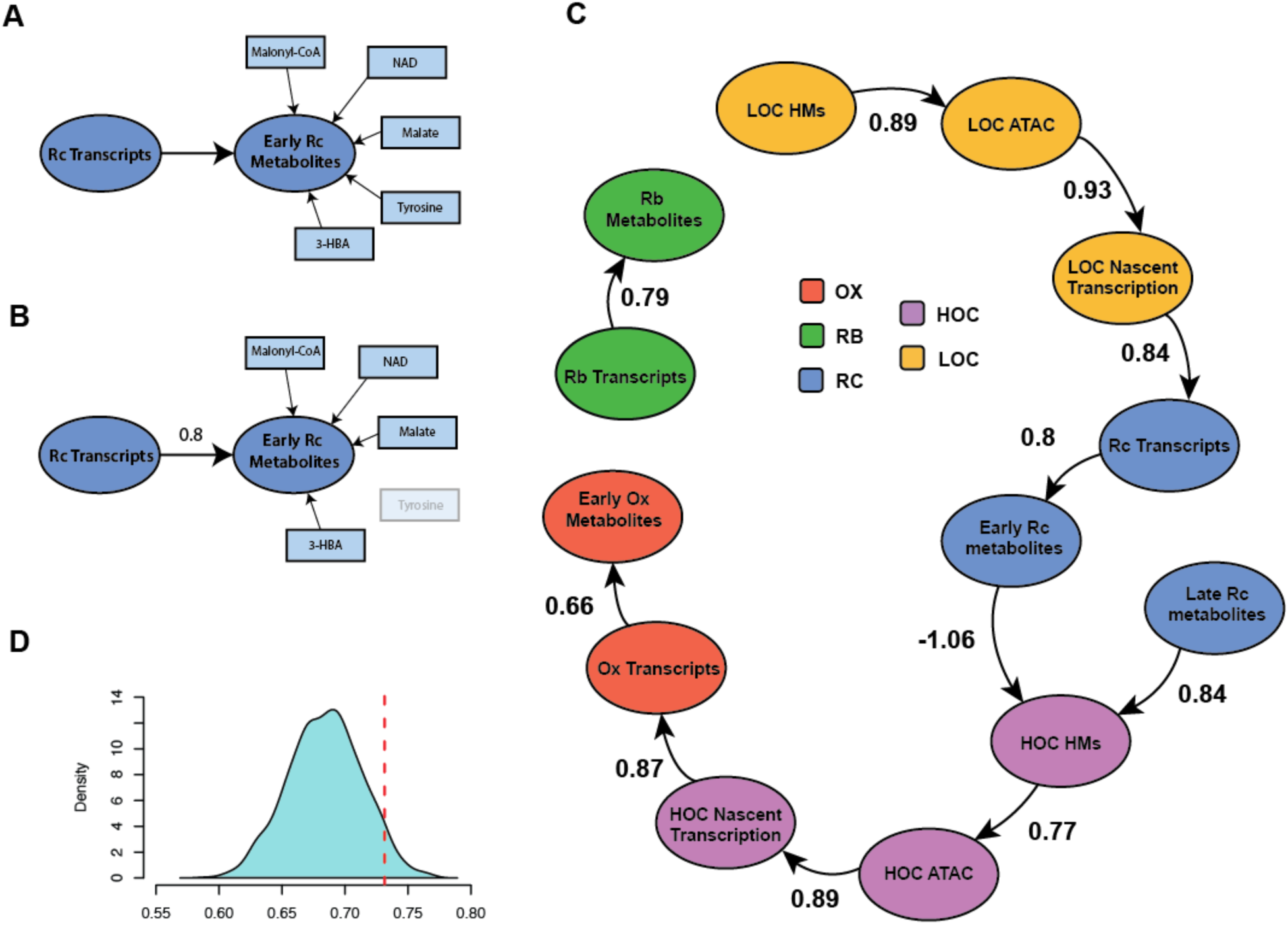
PLS-PM model recapitulates the molecular information that shapes the YMC. (A) Latent variable schematic, exemplifying two latent variables connected in the inner model (RC transcripts and Early RC metabolites) and the outer model indicators that construct the Early RC metabolites LV. (B) Same schematic as in (A) after fitting the model, the inner model coefficient is calculated and indicators that do not contribute to the model are discarded. (C) PLS path model after finalizing the iterative process of fitting the model, the numbers on the arrows represent the coefficient for the linear model fitted between each two latent variables. (D) Density of Goodness-of-Fit (GoF) values from 1000 random PLS-PMs. Our proposed model falls within the 95^th^ percentile of the GoF value distribution represented by the red dotted line.

Using this modelling framework, the PLS-PM algorithm fits the model to the observed data and quantifies the strength of each proposed regulatory relationship. We successfully characterized the connections among different cycling phases of metabolites, the two histone modifications combined into a single LV (see Methods), chromatin accessibility and both nascent and mature transcripts. Because the PLS-PM requires the model graph to be acyclic, and given that the RB phase is likely regulated post-transcriptionally, we removed the link between OX metabolites and RB transcripts (**Figure S3A**). This adjustment allowed us to focus our analyses on the LOC-HOC transition, which we hypothesize involves metabolic-epigenetic crosstalk.

Once the model was defined (**Figure S3A**) it underwent a two-step validation process: outer model validation and inner model validation. The outer model validation assessed how well the individual features contribute to their respective latent variable. This involved several steps. We first evaluated the contribution of each feature to its assigned latent variable, removing those that did not significantly contribute (**Figure 4B**). This refinement process was performed iteratively until the set of features for each latent variable stabilized (**Figure S3B**).

We also assessed the unidimensionality of each latent variable using the Chronbach’s alpha, a statistic that measures the internal consistency among the features contributing to a latent variable. During this process, several metabolites were eliminated from their latent variables. Analysis of their temporal profiles revealed that the model was sensitive enough to distinguish two types of metabolites within OX and RC phases: early-accumulating and late-accumulating (**Figure S3C**). This distinction was also observed in the initial PLS models (**Figure 2B-C**). Based on this finding, we refined the PLS-PM model to explicitly separate OX and RC metabolic phases into early and late accumulation phases (**Figures S3A and S3C**).

After the final outer model validation, all features exhibited loadings greater than 0.7 for their corresponding latent variable. All latent variables also showed high unidimensionality across multiple metrics. Specifically, the Chronbach’s alpha statistic was greater than 0.95 for all latent variables, except for the latent variable for late RC metabolites, which included only three features (**Table S3**). One latent variable (late OX metabolites) was excluded from the model due to an insufficient number of contributing features (**Figure 4C**).

Following validation of the outer model, the strength of the connections in the inner model was estimated by linear regression (**Figure 4C**). Any proposed connection yielding a non-significant regression coefficient was removed, and all coefficients were recalculated. In the final model, two connections (RB metabolites to LOC HMs and RC Transcripts to Late RC metabolites) were not significant (**Table S4**), and thus excluded (**Figure 4C**). The final model achieved a goodness of fit of 0.73, significantly higher than expected by chance (**Figure 4D**). Model significance was assessed by generating and fitting 1,000 random PLS-PM models (with randomized connections) using the same latent variables. Our model ranked within the top 5% in terms of goodness of fit (**Figure 4D**). Together, these results demonstrated that the proposed model robustly recapitulates the multilayered molecular organization of the YMC and provides a powerful framework for exploring how metabolites influence the transitions between HOC and LOC states.

### Metabolite accumulation connects the HOC and LOC states of the YMC

Latent variables (LVs) should consist of features that are highly correlated with the corresponding LV. To verify that features were correctly assigned, we examined their cross-loadings, which indicate whether a feature correlates more strongly with another LV than the one it was assigned to (**Figure S4**). Most features exhibited the highest loading within their assigned latent variable, confirming the internal consistency of the model. However, we identified a few cases where certain features had higher loadings in other LVs. For instance, some “HOC HMs” features correlated more with the “Ox Transcripts” LV (**Figure S4**). These cross-loadings likely reflect the intrinsic correlations among some LVs. Notably, we did not detect any cross-loadings between the HOC-LOC phases, further supporting the antagonistic nature of HOC and LOC states.

We next evaluated the connections among latent variables in the inner model (**Table 4**). The PLS-PM inner model effectively recapitulated the expected links between chromatin, transcription and gene transcripts. The significant regression coefficients in the connections reflect the high correlation among these molecular layers, revealing the tight molecular oscillations linking chromatin accessibility, histone modifications and gene transcription.

By contrast, capturing the associations between metabolites and their connected latent variables (transcripts and histone modifications) proved more challenging. The connection between Early RC Metabolites and HOC Histone Marks displayed a negative coefficient, whereas the links between Early RC Metabolites and RC Transcripts, as well as between RB Metabolites and LOC Histone Marks, were directly excluded from the final fitted model (**Table 4**, **Figure 4C**). These results suggests that metabolite LVs exhibit oscillatory dynamics that diverge from those of their associated molecular layers, likely due to the phase shifts in metabolite accumulation relative to transcript levels (**Figures 2A, 5A, S3B**). Such temporal offsets reduce the correlation strength and, consequently, affect the regression coefficients. Given that the transition between HOC and LOC phases coincides with periods of metabolite accumulation, shifting the cycle between the different energy states, we hypothesize that this accumulation plays a role in the epigenetic regulation of gene transcription. In the following section, we investigate which metabolites may introduce changes in chromatin state and explore why epigenetic regulation of transcription proceeds in two phases, while metabolites cycle in three phases.

**Figure 5.**
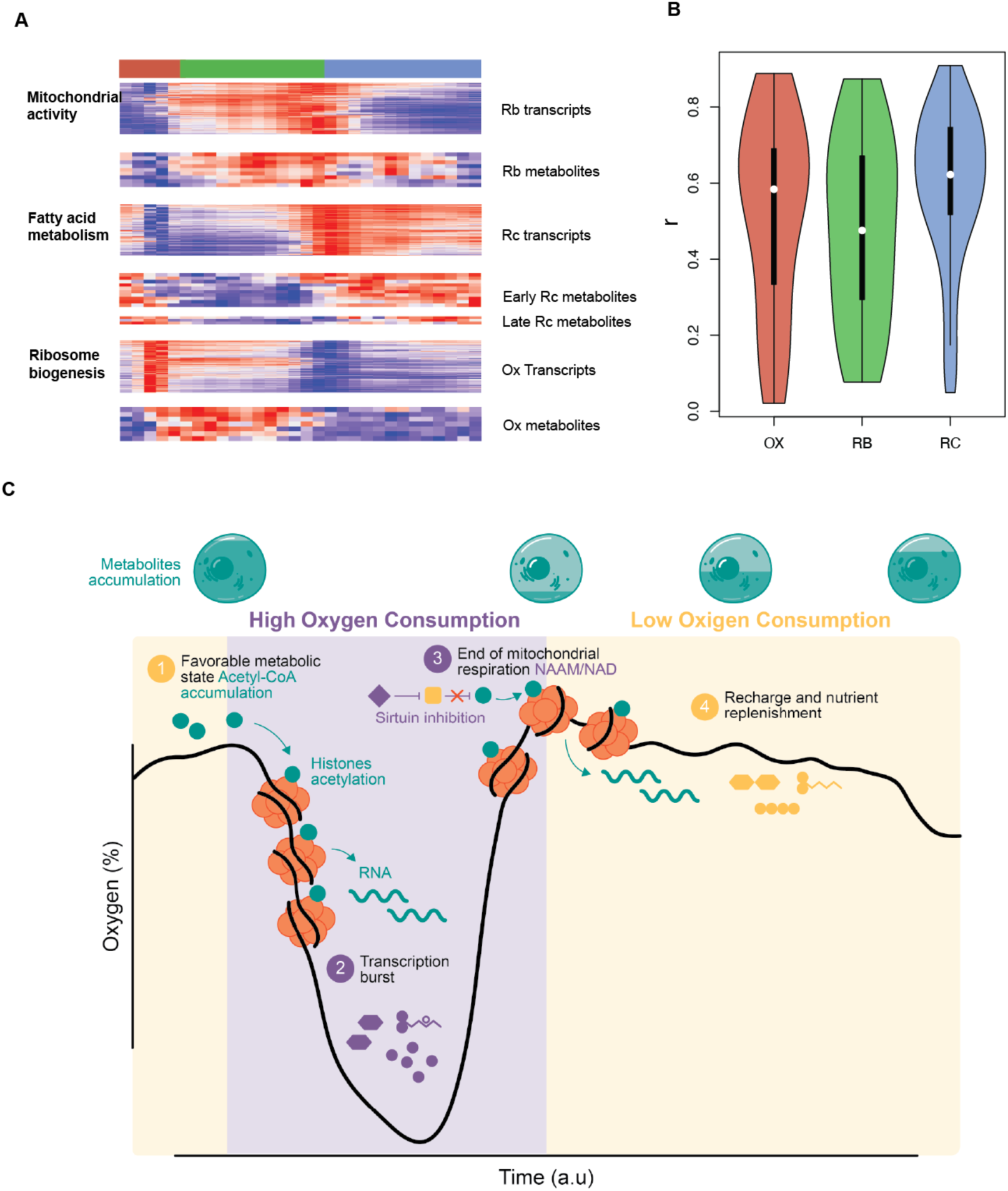
Key redox metabolites accumulate during HOC-LOC transitions to signal the cycle progression. (A) Heatmap of transcripts and metabolites that compose the indicated Latent Variables of the PLS-PM model (B) Per Gene correlation coefficient (r) resulting from correlating the peak quantification with their corresponding gene by proximity and separating in OX, RB and RC mature transcript phases (C) Proposed YMC biological model after analyzing the PLS-PM results, where HOC-LOC transitions are governed by metabolite accumulation that signal the transition to the subsequent phase of the YMC.

### Transition between HOC and LOC states

The PLS-PM model proved to be a powerful approach to establish the links among different omic layers across metabolic states and to identify elements that deviate from the correlation patterns proposed in the initial network. The iterative construction of latent variables ensured high sensitivity in selecting the features that best contributed to the model. Our results revealed two clearly distinct metabolic phases in the YMC, reflecting the physicochemical properties of the cycle. The high energy expenditure during the HOC phase drives the cells into a low-energy state, leading to the accumulation of NAM, NMPs and citrate (**Figure 2B**). NAD-derived metabolites are well-studied sirtuin cofactors^3,17,21^. In particular, NAM acts as a sirtuin inhibitor and is part of the metabolic sensor mechanism that regulate cellular activity. We propose that this process plays a central role in the HOC to LOC transition by modulating histone acetylation and thereby activating to the transcriptional program associated with LOC genes involved in fatty acid oxidation and peroxisome function^9^.

We next investigated the chromatin dynamics of the genes that remained in the outer model after fitting the PLS-PM model (**Figure 4C**), aiming to identify genes with chromatin states responsive to the HOC-LOC transition metabolites. Comparing ATAC profile across groups of genes, we found that the ATAC peaks showed stronger correlations with transcripts of metabolic genes such as *TDH3*, *CYB2* or *UGP1,* whose levels peak during the RC phase (**Figure 5B**). This indicates that chromatin dynamics are tightly regulated in the LOC phase, and that transcriptional activation of these “recovery” genes is more strongly influenced by the chromatin state than that of other genes in the cycle.

Our model supports a view of the Yeast Metabolic Cycle in which metabolite accumulation activates cellular sensors that drive transitions between HOC and LOC states (**Figure 5C**). Analysis of model loadings identified the most relevant features that govern each phase. Within the LOC phase LVs, including RC transcripts, we detected genes involved in characteristic processes of this stage, such as fatty acid oxidation (*POX1*, *POX2*) and trehalose metabolism (*TPK1*, *TPK2*, *ATH1*, *MAL11*, *NTH1*, *TPS2*). Metabolites produced by enzymatic activity in this phase were divided into two stages: an early set that contained 3-HBA, NTPs, aspartate, isoleucine, malonyl-CoA and the TCA intermediates fumarate and malate, all indicative of a high energy state (**Figure 5C**). During LOC phase, cells “recharge” their metabolic pool and the accumulation of these metabolites trigger the cell to enter HOC phase, partly by promoting histones acetylation to mediate gene expression^14^ (**Figure 5C**).

In contrast, LVs corresponding to HOC phase were enriched in genes associated with growth (*HXK2*) and nucleotide and transcription metabolism (*HTZ1*, *POL32*, *PNP1*). The HOC phase also featured strong expression of genes involved in amino-acid metabolism (*CYS3*, *SAM1*, *MET6*, *MET17*, *ARO10*) and the TCA cycle (*ACO1*, *IDH1*, *KGD1*), consistent with high mitochondrial activity previously reported^6^. By the end of the HOC phase, cells accumulate NAM, NAAM and NMPs, likely reflecting the consequences of elevated energy demands at this stage. We propose that the accumulation of these metabolites act as a signal for cells to enter the LOC phase by inhibiting sirtuin activity and altering the dynamics of the chromatin acetylation landscape (**Figure 5C**).

In summary, by applying PLS-PM, we successfully generated a multi-layered model that recapitulates the molecular dynamics of the YMC across epigenetic, transcriptional and metabolic levels. This model reflects the dominance of two principal molecular phases, HOC and LOC, and highlights specific metabolites that may mediate the transition between these states by regulating gene expression.

## Discussion

This study leverages the power of PLS-PM to construct a multi-layered model of molecular interactions revealing new regulatory associations within the Yeast Metabolic Cycle (YMC). We confirm that transcripts oscillate across the three canonical YMC phases (OX, RB and RC), while nascent transcription (NET-Seq), histone acetylation marks (H3K9ac and H3K18ac) and chromatin accessibility (ATAC-Seq), display two-phase dynamics corresponding to high and low oxygen consumption states (HOC and LOC), broadly reflecting cellular redox states (**Figure 2A, 5C**). Notably, our model indicates that chromatin-independent regulation of transcript cycling emerges during the RB phase, consistent with previous evidence of post-transcriptional control in this stage^8^. Although ATAC-seq offers lower resolution than the histone marks datasets, all chromatin-related layers exhibit two-phase oscillations closely correlated with nascent transcription (NET-seq). Chromatin remodeling was most prominent at genes producing mRNA transcripts clustering in OX and RC phases (**Figure 3D-G**). Moreover, joint modeling of transcript levels and ATAC-seq peaks revealed strong correlation during the RC phase, suggesting tighter chromatin regulation during this stage (**Figure 5B**).

Because metabolism acts both a consequence of protein activity and a regulator of gene expression, we reasoned that a multilayered integrative model encompassing all datasets and all conceptual YMC phases was required. PLS-PM proved particularly well suited for this purpose, as it allows the summarization of omics layers and phases as latent variables, assessment of the strength of their relationships, and data-driven refinements to the model structure. For instance, separating of early- and late-accumulating metabolites (**Figure 4C, S3A**) enhances sensitivity and helped identify metabolites with potential chromatin-modulation roles. We propose that low-energy metabolites (i.e. NAM, NMPs) accumulating at the HOC-LOC transition play a critical role in modulating chromatin structure and enabling the transcriptional reprogramming characteristic of early LOC^6^ (**Figure 5C**). This includes the activation of glucose-repressed genes (*GUT2* and *ADH2*), and genes involved in fatty acid degradation and trehalose metabolism. We suggest that the metabolites accumulating during late HOC repress sirtuin activity^21^ leading to increased H3K9ac and H3K18ac at the promoter of these genes^9^. The LOC phase, previously defined as a “Recharging” (RC) phase, is characterized by catabolic activity that replenishes the celĺs metabolic pools^7^, producing high-energy metabolites, such as NTPs, malonyl-CoA, and acetyl-CoA. Accumulation of these metabolites marks the LOC-to-HOC transition. Indeed, exogenous addition of acetyl-CoA to the culture is sufficient to trigger HOC onset and increase histone acetylation^14^ consistent with our observations of metabolites acting as signaling molecules.

While many metabolites exhibit two-phase associations during HOC-LOC transitions, their dynamics also align with the three-phase transcript oscillations. Integrated transcript-metabolite modelling revealed subsets of metabolites peaking in the OX, RB and RC phases (**Figure 2B** and **C**), offering new insights into the ultradian rhythms regulation characteristic of the YMC^1^. These three-phase associations may arise from post-translational modifications of metabolic enzymes, such as phosphorylation, that fine-tune metabolic flux. Such regulation was previously demonstrated for Nth1, involved in trehalose degradation and for Rps6, associated with TORC1 activity^22,23^. This contrasts with our proposed two-phase cycling metabolites, which are tied to redox balance and act directly on chromatin architecture.

Based on our multi-layer model, we propose that chromatin modifications serve as a two-phase metabolic readout, while modulation of enzyme activity constitutes a three-phase readout, with transitions driven by the accumulation of high- and low-energy metabolites. These transitions vary depending on the molecular layers examined (e.g., NET-seq versus RNA-seq). Our previous work suggests that transcriptional changes alone – whether at nascent or steady-state levels – are insufficient to drive YMC phase transitions, as the proteome remains relatively stable ^8^, which points to a role in replenishing total protein pools diluted by growth and turnover rather than acting as the primary oscillator^8^. Instead, fluctuations in high- and low-energy metabolites likely exert their regulatory effects through post translational modifications, acting on chromatin in the nucleus and on metabolic enzymes in the cytoplasm.

Finally, the weaker model fit for certain metabolites may reflect rapid turnover during energy state transitions. Metabolites synthesized at high rates may be rapidly consumed to meet immediate cellular demands, resulting in transient accumulation at phase transition. Additionally, elevated histone acetylation could act as a temporal sink for high-energy metabolites, indirectly promoting transcriptional changes^24^, an effect potentially mirrored by analogous enzyme phosphorylation cycles.

In summary, our study demonstrates how a complex ultradian system such as the YMC can be dissected by integrative multi-omics modeling to uncover the coordinated regulation of its molecular layers. These findings reveal that metabolites, chromatin, and transcriptional programs function as interdependent switches coordinated with progression through redox cycles, providing a conceptual framework applicable to oscillatory systems across biological contexts.

## Methods

### Fermenter conditions

Yeast strain used for this study is CENPK113-7D rpb3-FLAG, cells were cultured in fermenter BioFlo320 (2L vessel, 1.1 L culture volume – New Brunswick), cultures were grown in YMC-YE media (pH 3.5 – ammonium sulphate 5 g/l, potassium dihydrogen monophosphate 2 g/l, magnesium sulphate 0.5 g/l, calcium chloride 0.1 g/l, yeast extract 1 g/l (Difco), glucose 10 g/l, sulphuric acid 0.035 %, antifoam-204 0.05 % (Sigma Aldrich), iron sulphate 20 mg/l, zinc sulphate 10 mg/l, manganese chloride 1 mg/l, copper sulphate 10 mg/l). Fermenter runs had an aeration rate on 1 l/min and agitation rate of 1000 rpm. Runs temperature was set to 30°C and pH was constant at 3.5 through the addition of 0.25 NaOH.

Starter cultures (10 mL) were grown to saturation overnight at 30°C, starved for 6 h, then used to initiate continuous cultures (1.5 mL/min flow rate). Cultures were stabilized for ≥24 h before sampling. Yeast metabolic cycles (YMCs) were synchronized as described^7^, with sampling adjusted to obtain balanced time points across cycle phases^6^.

### RNA-seq

RNA-seq data (GSE52339) from 16 YMC time points (Illumina HiSeq 2000, 50 bp single-end) were processed as described^6^, with depth normalization, log transformation, and gene-wise centering.

### ChIP-seq processing

We used two histone modification ChIP-seq experiment for H3K9ac (GSE52339) and H3K18ac (GSE118889), each sampled in 16 time-points evenly distributed across the YMC. Raw reads were quality-controlled using FastQC and trimmed with Trimmomatic^25^ (minimum quality score 30, sliding window 5, minimum length 28 bp). Reads were aligned to the *S. cerevisiae* Ensembl R91 reference genome using Bowtie2. We quantified signal in 300 bp regions upstream of transcription start sites using bedtools genomecov and normalized to H3 ChIP-seq controls, followed by log transformation and centering.

### NET-seq

Nascent transcript sequencing count data was obtained from GSE138023. The dataset was evenly sampled in 12 time-points across the cycle. We applied TMM normalization, log transformation and centering.

### ATAC-seq

Yeast cells for ATAC-Seq were evenly sampled in 18 timepoints across the cycle in two batches. Cells were fixed in 1% formaldehyde for 5min at room temperature, glycine was then added to a final concentration of 125mM and incubated for 5min. The equivalent to 30 million cells were washed with water, resuspended in Zymolyase preincubation buffer (250μL 14.5M β-ME, 27.8μL 0.5M EDTA pH 8 and complete to 5mL with Sorbitol 1M) and kept at 30° for 30 min. Cells were washed in sorbitol, pellet was resuspended in 500 µL of Zymolyase buffer (3.5μL 14.5M of B-ME, complete to 10mL with Sorbitol 1M, resuspend Zymolyase to a final concentration of 10mg/mL) and incubated 30° for 12 min to digest yeast cell wall. Cells were washed gently 2x in sorbitol and 2x in PBS to eliminate β-ME remains, and then resuspended in lysis buffer (10mM Tris-HCl pH 7.4, 10mM NaCl, 3mM MgCl2, 0.1% IGEPAL CA-630) and centrifuged at 10000 rpm for 10 min. Supernatant was discarded, pellet included isolated nuclei that were resuspended in 50µL of transposition mix and incubated at 37° of 30 min. When transposition reaction finishes, 10µL of STOP buffer (85% H2O, 5% SDS, 10% 0.5M EDTA) was added to the tubes, DNA was de-crosslinked using 7µL of NaCl 5M and incubating for 3h at 65°. Then 1µL of proteinase K was added and incubated at 65° overnight.

DNA was purified using Qiagen MinElute PCR purification Kit, transposed DNA was eluted in 11µL of elution buffer. DNA was PCR amplified using Nextera DNA Library Preparation Kit and Nextera Index Kit and purified using AmpureXP cleanup for a final insert size of 150-180 bp. Quality of the library size was assessed using Qubit, libraries were sequenced with Illumina NextSeq 500 with NextSeq 500/550 High Output Kit v2.5 (75 Cycles). ATAC-Seq data quality was assessed with Fastqc (Andrews, 2010), and adapter and quality trimmed with Trimmomatic^26^ using a minimum quality of 30, a sliding window of 5 and a minimum length of 28 bp. Data was aligned to the Ensembl *S. cerevisiae* reference genome (release 91) using Bowtie mapper^27^. We quantified promoter accessibility by using read counts of 300bp upstream of TSS. Six time-points were low-quality filtered (Figure S1D), which resulted in 12 time-points ATAC measurements (Figure 1A).

Peaks were identified with MACS^28^ merging the alignment files from time-points in the Low Oxygen Consumption (which includes time points 8 to 12) and High Oxygen consumption (which includes time points 1 to 7) phases. TF binding site motifs were identified using the Homer software, by using the findmotifsgenome.pl function and using the JASPAR database as reference^29^.

Peaks were tested for differential expression between HOC and LOC timepoints using DESeq and using the different timepoints of each phase as replicates. Heatmaps were constructed using ComplexHeatmap package in R.

### Metabolomics

For metabolomics sampling, yeast cells were harvested from BioFlo fermenter, 9 tubes with 20×10^6^ cells per sample. Cells were pelleted, supernatant removed and snap-frozen with liquid nitrogen. A total of 105 samples were extracted and evenly distributed in 5 cycles of the YMC, which resulted in 21 timepoints distributed in 5 cycles.

Cell pellets were lyophilized to dryness overnight. The lyophilized pellet was homogenized in 50:50 acetonitrile/water with 0.3 % formic acid for Oas and Aas, 5% TCA for acetyl and malonyl CoA, 0.5M PCA for oxidized nucleotides and 50:50 methanol/NaOH for reduced nucleotides using the Precellys beadbased homogenizing system. Precellys tough micro-organism lysing kit (VK-05) was used to burst the yeast cells. An aliquot of homogenate for the required assay was aliquoted and immediately stored at −80°C.

Oxidized nucleotides were separated on a Thermo Scientific Hypercarb column (3 µm x 50 mm x 2.1 mm) using a Thermo Dionex Ultimate 3000. Nucleotides reduced were separated on waters HSST3 1.8 µm x 2.1 mm x 150 mm column. Organic acids were separated on Waters Acquity UPLC BEH C18 2.1 mm x 100 mm, 1.7 µm column. Amino acids were separated on a 2.1 x 100 mm, 1.7 µm Waters AccQTag column. CoAs were separated by waters Acquity UPLC BEH C18 1.7 µm x 2.1 mm x 50 mm column. A total of 63 metabolites were separated with the chromatographies and quantified using a Thermo Scientific Quantiva Triple Quadrupole mass spectrometer. The mass spectrometer was operated in positive ion mode using electrospray ionization. The raw data was processed using Xcalibur 3.0. Data was corrected for protein abundance and median normalized. The most extreme value of each metabolite was discarded, and the resulting matrix was centered.

### Differential expression and clustering

The temporal profile of each omic feature was modeled with the maSigPro R package^30^ using a maximum polynomial degree of 3. Feature significantly changing across time (FDR adjusted p-value < 0.05) were additionally required to have a minimum R^2^ value of 0.6 for gene expression, histone modifications and NET-Seq, and 0.4 for ATAC-Seq and metabolomics.

For each omic, these differentially expressed (DE) features were scaled and clustered with the k-means algorithm. We tested different number of optimal clusters using the k-means algorithm and different clustering quality metrics (“kl”, “ch”, “16ubert16n”, “cindex”, “db”, “silhouette”, “duda”, “pseudot2”, “beale”, “ratkowsky”, “ball”, “ptbiserial”, “gap”, “frey”, “mcclain”, “dunn”, “16ubert”, “sdindex”, “dindex”, “sdbw”), implemented in the nbclust R package^31^. We then ranked the number of clusters that were appointed by each clustering index metric to decide the number of clusters that were going to be used to model each omic.

### Matching omics datasets

The different omics datasets combined in this study were extracted from different YMC experimental runs and had slightly different positions at the cycle (Figure 1A). We aligned all omics datasets to 30 common time points using spline regression (11-21 knots depending on dataset) taking oxygen consumption profiles as temporal references.

### Partial Least Squares Regression

We performed PLS regression to model gene expression (**Y**), measured with either RNA-se or NET-seq, as a function of metabolite measurements (**X**), using the equation **Y** = **XB** + **E**, where **B** is the matrix containing the regression coefficients and **E** the residuals^13^. **X** and **Y** matrices were centered and scaled for this analysis. We optimized the number of components by optimizing R^2^ and Q^2^ and two components were selected in both cases.

### PLS Path Modeling

PLS Path Modeling (PLS-PM) is a component-based method for estimating Structural Equation Models (SEM)^20^. The PLS-PM methodology estimates and tests a set of relationships between variables that can either be measured or unobserved (latent variables), which can be represented in a theoretical network (path model) that is considered the starting hypothesis of the PLS-PM model. We constructed a path model with 15 latent variables (LV) representing different omics measurements in different phases across the YMC. The path model consists of two sub-models: the outer or measurement model and the inner or structural model. The outer model connects the observed data (indicators or manifest variables) with their associated LV *Y_j_*, which is calculated as a weighted sum of its *K_j_* indicators:

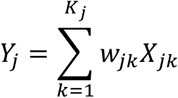

Where *X_jk_* is the indicator variable *k* associated to *LV Y_j_*, and the coefficient *w_jk_* is the corresponding weight (loading). The inner model is the one connecting the LVs, and these connections are estimated by multiple linear regressions:

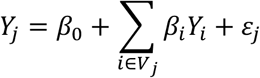

Where *V_j_* is the set of LVs that potentially predict Y_j_, β_l_ are the path coefficients and χ_j_ is the error term.

The clusters of differential features for any given omic were used to assign the indicators (outer model) of each LV in the inner model. The outer model was validated by measuring the indicators loadings (>0.7) and their unidimensionallity (Cronbach’s α >0.7):

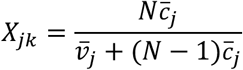

Where *N* is equal to the number of indicators in the LV *j*, 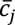 is the average covariance among the indicators of the LV and 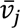 is the average variance of each individual indicator. The functions to calculate an LV unidimensionality were extracted from the plspm R package in CRAN^19^ and modified to allow more observations than variables. Indicators that did not satisfy these requirements were removed from the model.

The outer model validation suggested that OX and RC metabolites should be subdivided into two further LVs, since two separate subsets of metabolites in these phases were observed.

The inner model was constructed by connecting 14 latent variables based on their expected biological relationship. By fitting the model, we test whether these connections are supported by the data. Each of the 14 LVs corresponds to a given omic and phase, with two exceptions: the division of early and late metabolites for OX and RC phases and the combination of the two histone marks (H3K9ac and H3K18ac) into the same LV due to the high correlation between them (Figure S2B).

The inner model regression showed the unfeasibility of including the late OX metabolites’ LV in the inner model, since all the relationships connecting the LV showed non-significant coefficients and the late Ox metabolites LV got disconnected and removed from the model. Therefore, the final fitted PLS-PM model contained 13 LVs.

After model fitting, we also studied traitor indicators, which refer to indicators that have a higher correlation with a latent variable other than their own.

The path coefficients of the relationships that connect the LVs in the inner model determine whether a connection is supported by the data or not. These connections are estimated by linear regression models and evaluated using their R^2^ coefficient. A bootstrapping procedure was applied to compute the confidence intervals for the regression coefficients of each relationship^20^.

The overall performance of the model was evaluated using its Goodness-of-Fit (GoF). In order to check that our model was significantly more meaningful than a random model with the same LVs, we simulated 1000 random models with the same latent variables and randomized their relationships. We obtained the GoF values of each random model and created a null distribution for the GoF with these values. Our model’s GoF was among the top 5% random models, which proved its reliability and robustness.

## Data and code availability

Gene expression and histone modification data were retrieved from the Gene Expression Omnibus (GEO) repository (accession number GSE52339 and GSE118889). Raw data for ATAC-seq dataset can be obtained in the GEO under the accession number GSE146350. Metabolomics data is available on the metabolomics workbench under the accession number ST001350. Original code with data post-processing and analysis can be found in GitHub (https://github.com/ConesaLab/YMC_plspm) and is publicly available as of the date of publication.

## Acknowledgements

This work was supported by the Marie Skłodowska-Curie Grant Agreement No 675610, as part of the ChroMe (Chromatin and Metabolism) International Training Network. The metabolomics data were obtained through a pilot grant from the National Institute of Health’s (NIH) Common Fund Metabolomics Program (Grant number: U24DK097209) and were processed at the core facility for targeted metabolomics at Sanford-Burnham, led by Steven Gardell. Further funding was provided by the BBSRC Project Grant BB/S009035/1, “Using Mathematical Modelling to Deconstruct Transcription.” The authors thank Anna Lamstaes for assistance with the ATAC-seq library preparation and Jack Feltham for their contribution to setting up the fermenter.

## Author contributions

Salvador Casani-Galdon: Data extraction; Data curation; Data analysis; Figure generation; Writing-review and Editing. Shidong Xi: Data extraction. Jane Mellor: Conceptualization; Supervision; Writing-Review and Editing. Sonia Tarazona: Conceptualization; Supervision; Methodology; Writing-Review and Editing. Ana Conesa: Conceptualization; Supervision; Methodology; Writing-Review and Editing.

---

**Appendix Figure S1.**
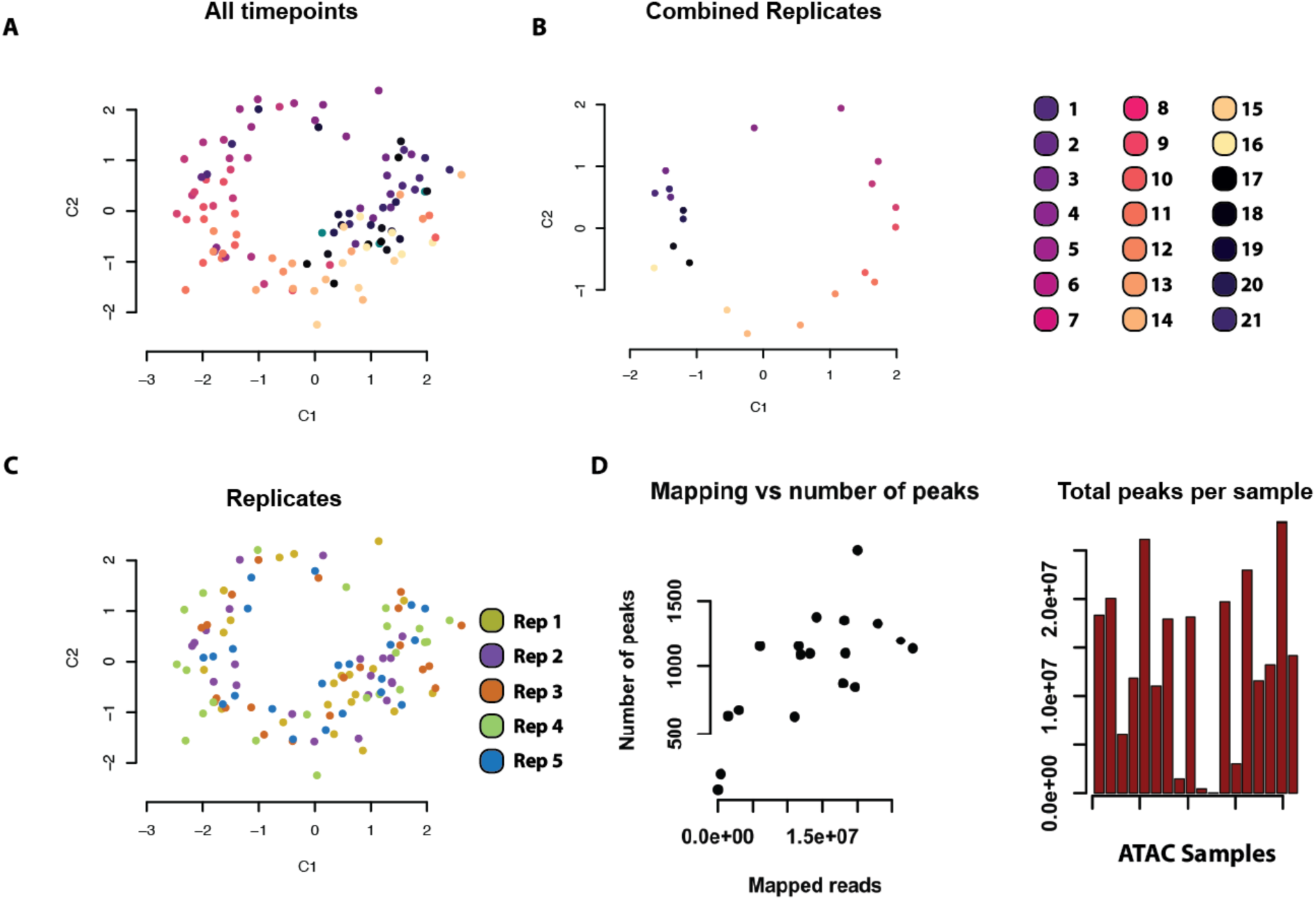
Quality control of metabolomics and ATAC-seq datasets. (A) PCA score plot of metabolites colored by time point (B) PCA score plots of collapsed time-points (C) PCA score plot of metabolites colored by replicate (D) ATAC QC: Number of ATAC peaks by number of reads per sample (left) and total number of reads in peak (right)

**Appendix Figure S2.**
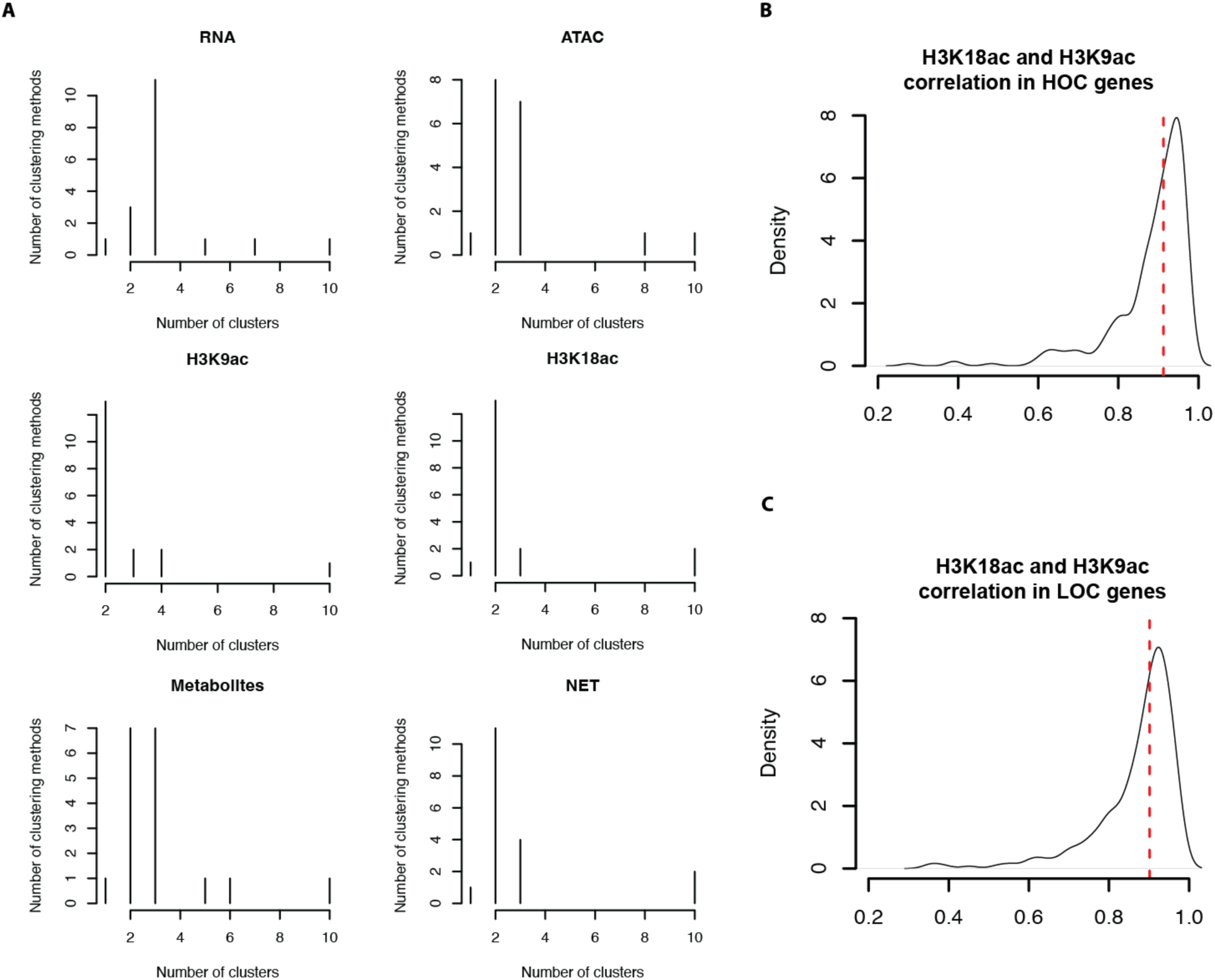
Clustering of the datasets. (A) Number of quality metrics that support a given number of clusters. (B) and (C) Density plot of the correlation of HOC (B) and LOC (C) gene measurements between H3K9ac and H3K18ac signals, the red dotted line points to the median, which is 0.913 and 0.901 for HOC and LOC, respectively.

**Appendix Figure S3.**
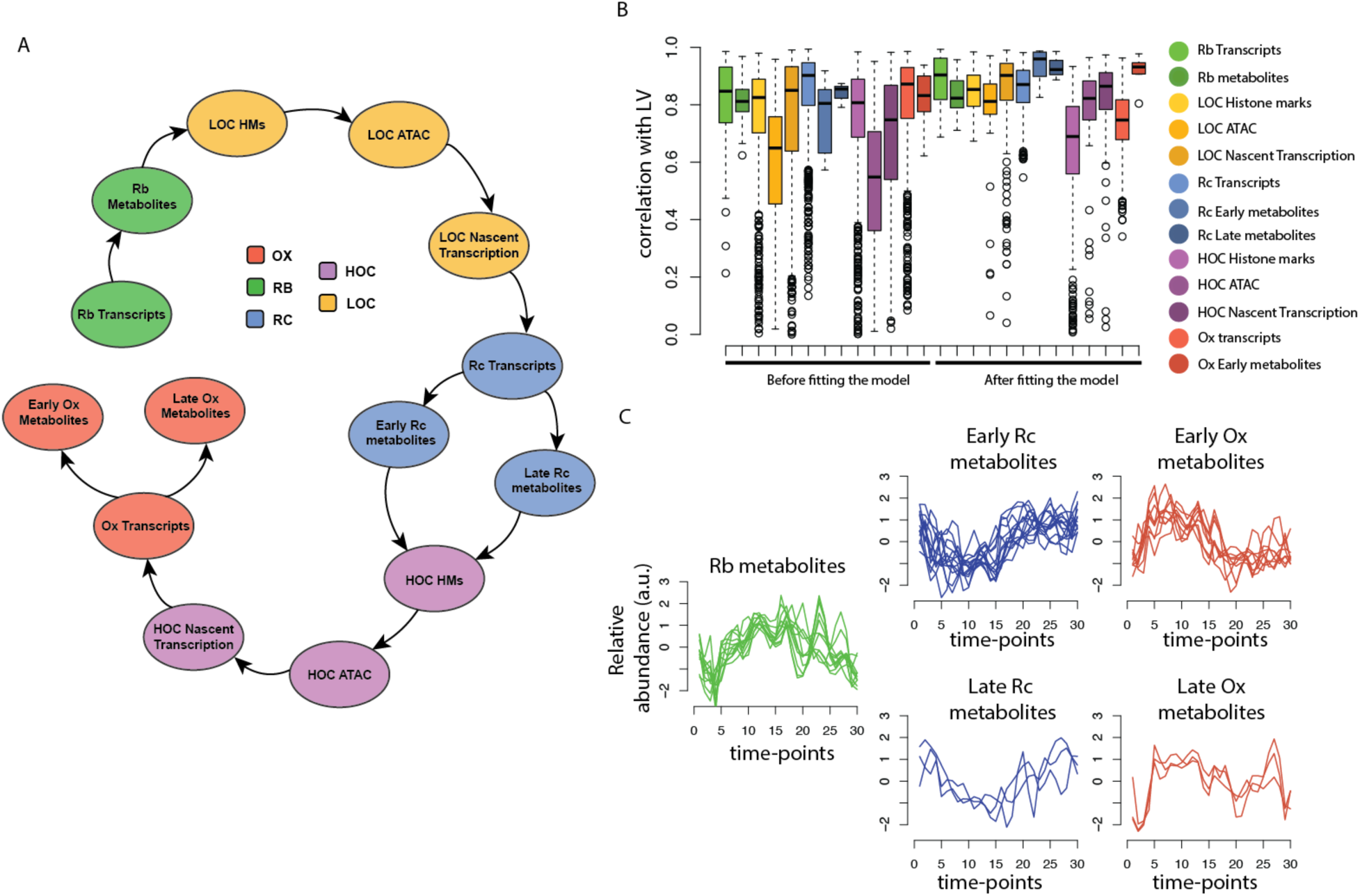
PLS-PM characterizes two metabolic subtypes for OX and RC phases. (A) Proposed PLS-PM inner model (B) Correlation of the elements in each latent variable before and after fitting the PLSPM model. (C) Metabolic profiles of the different groups, emphasizing in early and late metabolites

**Appendix Figure S4.**
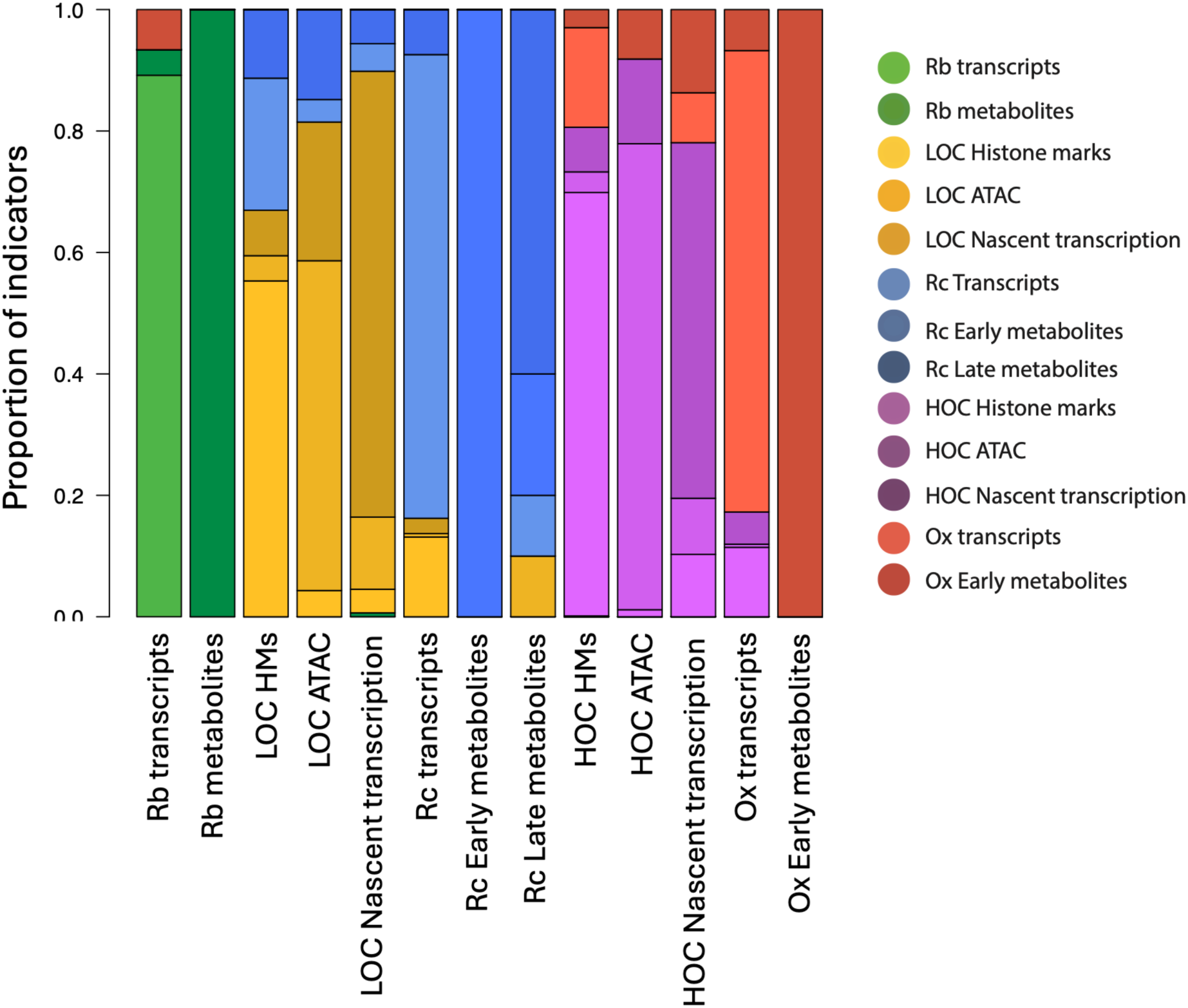
Cross-loading values of features in the latent variables.

Appendix Table S1. List of Metabolites profiled in this analysis.

Appendix Table S2. Number of features per phase in each molecular type.

Appendix Table S3. Unidimensionality measurements used for outer model validation.

Appendix Table S4. Inner model coefficients.

## References

1. Mellor, J. The molecular basis of metabolic cycles and their relationship to circadian rhythms. Nature Structural and Molecular Biology vol. 23 1035–1044 Preprint at 10.1038/nsmb.3311 (2016).

2. Pinu, F. R. et al. Systems biology and multi-omics integration: Viewpoints from the metabolomics research community. Metabolites 9, (2019).

3. Etchegaray, J. P. & Mostoslavsky, R. Interplay between Metabolism and Epigenetics: A Nuclear Adaptation to Environmental Changes. Molecular Cell vol. 62 695–711 Preprint at 10.1016/j.molcel.2016.05.029 (2016).

4. Kinnaird, A., Zhao, S., Wellen, K. E. & Michelakis, E. D. Metabolic control of epigenetics in cancer. Nature Reviews Cancer vol. 16 694–707 Preprint at 10.1038/nrc.2016.82 (2016).

5. Koronowski, K. B. & Sassone-Corsi, P. Communicating clocks shape circadian homeostasis. Science vol. 371 Preprint at 10.1126/science.abd0951 (2021).

6. Kuang, Z. et al. High-temporal-resolution view of transcription and chromatin states across distinct metabolic states in budding yeast. Nat Struct Mol Biol 21, 854–863 (2014).

7. Tu, B. P., Kudlicki, A., Rowicka, M. & Mcknight, S. L. Logic of the Yeast Metabolic Cycle: Temporal Compartmentalization of Cellular Processes. https://www.science.org.

8. Feltham, J. E. et al. Transcriptional changes are regulated by metabolic pathway dynamics but decoupled from protein levels. Preprint at 10.1101/833921 (2019).

9. Sánchez-Gaya, V. et al. Elucidating the Role of Chromatin State and Transcription Factors on the Regulation of the Yeast Metabolic Cycle: A Multi-Omic Integrative Approach. Front Genet 9, (2018).

10. Cesur, M. F., Çakır, T. & Pir, P. Genome-Wide Analysis of Yeast Metabolic Cycle through Metabolic Network Models Reveals Superiority of Integrated ATAC-seq Data over RNA-seq Data. mSystems 7, (2022).

11. Tu, B. P., et al. Cyclic Changes in Metabolic State during the Life of a Yeast Cell. vol. 104 (2007).

12. Buenrostro, J. D., Wu, B., Chang, H. Y. & Greenleaf, W. J. ATAC-seq: A method for assaying chromatin accessibility genome-wide. Curr Protoc Mol Biol 2015, 21.29.1–21.29.9 (2015).

13. Geladi, P. & Alski, B. R. K. PARTIAL LEAST-SQUARES REGRESSION: A TUTORIAL. Analytica zyxwvutsrqponmlkjihgfedcbaZYXWVUTSRQPONMLKJIHGFEDCBA Chimica Acta vol. 186 (1986).

14. Cai, L., Sutter, B. M., Li, B. & Tu, B. P. Acetyl-CoA Induces Cell Growth and Proliferation by Promoting the Acetylation of Histones at Growth Genes. Mol Cell 42, 426–437 (2011).

15. Burnetti, A. J., Aydin, M. & Buchler, N. E. Cell cycle start is coupled to entry into the yeast metabolic cycle across diverse strains and growth rates. Mol Biol Cell 27, 64–74 (2016).

16. Rao, A. R. & Pellegrini, M. Regulation of the yeast metabolic cycle by transcription factors with periodic activities. BMC Syst Biol 5, (2011).

17. Covarrubias, A. J., Perrone, R., Grozio, A. & Verdin, E. NAD+ metabolism and its roles in cellular processes during ageing. Nature Reviews Molecular Cell Biology vol. 22 119–141 Preprint at 10.1038/s41580-020-00313-x (2021).

18. Mentch, S. J. et al. Histone Methylation Dynamics and Gene Regulation Occur through the Sensing of One-Carbon Metabolism. Cell Metab 22, 861–873 (2015).

19. Tenenhaus, M., Vinzi, V. E., Chatelin, Y. M. & Lauro, C. PLS path modeling. Comput Stat Data Anal 48, 159–205 (2005).

20. Sanchez, G. PLS Path Modeling with R. www.gastonsanchez.com.

21. Anderson, K. A., Madsen, A. S., Olsen, C. A. & Hirschey, M. D. Metabolic control by sirtuins and other enzymes that sense NAD+, NADH, or their ratio. Biochimica et Biophysica Acta - Bioenergetics vol. 1858 991–998 Preprint at 10.1016/j.bbabio.2017.09.005 (2017).

22. Dengler, L., Örd, M., Schwab, L. M., Loog, M. & Ewald, J. C. Regulation of trehalase activity by multi-site phosphorylation and 14-3-3 interaction. Sci Rep 11, (2021).

23. Nakashima, A., Sato, T. & Tamanoi, F. Fission yeast TORC1 regulates phosphorylation of ribosomal S6 proteins in response to nutrients and its activity is inhibited by rapamycin. J Cell Sci 123, 777–786 (2010).

24. Charidemou, E. & Kirmizis, A. A two-way relationship between histone acetylation and metabolism. Trends in Biochemical Sciences vol. 49 1046–1062 Preprint at 10.1016/j.tibs.2024.10.005 (2024).

25. Quinlan, A. R. & Hall, I. M. BEDTools: A flexible suite of utilities for comparing genomic features. Bioinformatics 26, 841–842 (2010).

26. Bolger, A. M., Lohse, M. & Usadel, B. Trimmomatic: A flexible trimmer for Illumina sequence data. Bioinformatics 30, 2114–2120 (2014).

27. Langmead, B. & Salzberg, S. L. Fast gapped-read alignment with Bowtie 2. Nat Methods 9, 357–359 (2012).

28. Feng, J., Liu, T., Qin, B., Zhang, Y. & Liu, X. S. Identifying ChIP-seq enrichment using MACS. Nat Protoc 7, 1728–1740 (2012).

29. Sandelin, A., Alkema, W., Engström, P., Wasserman, W. W. & Lenhard, B. JASPAR: An open-access database for eukaryotic transcription factor binding profiles. Nucleic Acids Res 32, (2004).

30. Nueda, M. J., Tarazona, S. & Conesa, A. Next maSigPro: Updating maSigPro bioconductor package for RNA-seq time series. Bioinformatics 30, 2598–2602 (2014).

31. Charrad, M., Ghazzali, N., Laval, U. & Niknafs, A. NbClust: An R Package for Determining the Relevant Number of Clusters in a Data Set Véronique Boiteau. JSS Journal of Statistical vol. 61 http://www.jstatsoft.org/ (2014).

